# Natural product-based putative efflux inhibitors restore bedaquiline susceptibility in a drug-resistant *Mycobacterium tuberculosis* mutant

**DOI:** 10.64898/2026.06.08.730784

**Authors:** Robi Chacha, Maes Valerie, Mathew P. Ngugi, Edwin K. Murungi, Dirk A. Lamprecht, Elizabeth M. Kigondu

## Abstract

Tuberculosis (TB) caused by *Mycobacterium tuberculosis* (*Mtb*) remains a potent threat to global public health. Moreover, the alarming surge in the number of multidrug-resistant (MDR) and extensively drug-resistant (XDR) *Mtb* strains will continue to imperil TB control efforts. Thus, the discovery of new TB agents with novel modes of action or resistance-reversing therapeutics is a pressing priority. In this study, we report the generation of spontaneous *Mtb* mutants exhibiting bedaquiline (BDQ) resistance and the subsequent evaluation of natural product-derived efflux inhibitors (EIs) that restored the antimicrobial efficacy of BDQ against the mutants. BDQ-resistant mutants were successfully isolated, and colonies were observed on agar plates with concentrations up to 100x the minimum inhibitory concentration (MIC). Upon screening against BDQ, the resistant strains exhibited MIC values ranging from 0.098 μM to 3.136 μM, corresponding to 1–32-fold increases relative to the wild type, with higher resistance observed on 50x- and 100x-selection plates. Genetic analysis identified point mutations and frameshifts in key resistance-related genes, including *Rv0678*, *pepQ*, and *atpE*. Notably, combining BDQ with EIs such as berberine (BER), reserpine (RES), piperine (PIP), and lyoniresinol (LYO) remarkably lowered the MIC in the selected mutant strain. Synergistic effects were observed for BDQ+BER (FICI = 0.188; 16-fold MIC reduction) and BDQ+RES (FICI = 0.37; 8-fold MIC reduction). For BDQ+LYO, the FICI could not be calculated because LYO did not have an MIC at the highest concentration tested; however, this combination produced the strongest effect, restoring susceptibility with a 64-fold MIC reduction and exhibiting bactericidal activity. These results highlight the role of efflux pumps in BDQ resistance and support the use of natural product-derived EIs as potential supplementary therapies against drug-resistant *Mtb*.

## Introduction

Tuberculosis (TB), caused by *Mycobacterium tuberculosis* (*Mtb*), remains a potent global public health threat with approximately 10.7 million new cases and 1.23 million deaths reported in 2024 (World Health Organization (WHO), 2025). The burden of the disease is disproportionately high in resource-constrained regions of Africa and Southeast Asia and may increasingly worsen due to the surging number of multidrug-resistant (MDR) and extensively drug-resistant (XDR) *Mtb* strains. Recent studies indicate that approximately 500,000 new cases of drug-resistant TB are reported yearly, largely in Africa and Southeast Asia regions (Goletti *et al*., 2025). Resistance to BDQ, a cornerstone drug for the treatment of MDR and XDR TB, is a particularly concerning development (Saeed *et al*., 2022). By targeting the *atpE* gene encoded c-subunit of *Mtb* ATP synthase, BDQ inhibits the critical mycobacterial ATP synthesis (Xu *et al*., 2023). Given that drug extrusion is the primary mechanism underpinning BDQ resistance (Kanji *et al*., 2019), targeting *Mtb* efflux pumps (EPs) is a viable strategy for developing novel TB therapeutics (Yang *et al*., 2020).

Of the more than 50 EPs identified in *Mtb*, MmpS5-MmpL5, Rv1258c, and Rv1819c have been strongly implicated in MDR and XDR TB infections (Kanji *et al*., 2019; Rodrigues *et al*., 2020). MmpS5-MmpL5 mediates efflux of BDQ, clofazimine (CFZ), azoles, tetracyclines and several dugs in clinical development including TBAJ-876, PBTZ-169 PBTZ-169 and OPC-167832 (Fountain *et al*., 2025). On the other hand, Rv1258c extrudes rifampicin (RIF), isoniazid (INH), pyrazinamide (PZA), ethambutol (EMB) and some aminoglycosides (Laws *et al*., 2022). Mutations in Rv0678, atpE and pepQ genes are linked to the over-expression of MmpS5-MmpL5 (Almeida *et al*., 2016; Andries *et al*., 2005; Hartkoorn *et al*., 2014). Natural products (NPs) have been shown to possess the ability to block efflux (Ramalingam *et al*., 2024; Seukep *et al*., 2020) and may thus be explored as probable EIs. For instance, reserpine has been demonstrated to halt tetracycline efflux in *Bacillus subtilis* (Shaheen *et al*., 2019) while piperine has been shown to interfere with *Mtb* Rv1258c (Sharma *et al*., 2010). Besides, berberine has been reported to derange MdfA in *Escherichia coli* (Li & Ge, 2023; Morita *et al*., 2016). Experimental evidence demonstrating the capacity of natural products derived EIs to reverse BDQ resistance in *Mtb* remains scanty. This study evaluated the potential of several NP derived compounds to restore *Mtb* susceptibility to BDQ potentially through efflux pump inhibition in an Rv0678 mutant.

## Materials and methods

### Selection of putative efflux inhibitors

Candidate EIs were identified through an extensive review of literature on NP derived compounds reported to enhance the antimicrobial activity of standard drugs. Relevant studies describing phytochemical origins and synergistic effects of EIs were evaluated, and the corresponding source plant species were documented (Mokhber-Dezfuli *et al*., 2014; Puk & Guz, 2022; Seukep *et al*., 2020). Subsequently, phytochemicals isolated from the identified plant species were ascertained, and potential EIs were parsed based on structural similarity to compounds reported to inhibit efflux. To broaden the number of potential EIs, a search of chemical and structural analogs to the identified compounds in public databases, including ZINC, PubChem, and ChemSpider, was performed. Specifically, to constrain the search to relevant candidates, database queries utilized structural similarity and substructure search parameters. Following physicochemical profiling, the antimicrobial activity of selected compounds was determined using *in vitro* assays.

### Drugs and reagents

Berberine chloride (BER, Cat. No. B3251), reserpine (RES, Cat. No. R0875), piperine (PIP, Cat. No. P49007), bedaquiline fumarate (BDQ, Cat. No. SBR00060-10MG), doxycycline hyclate (DOX, Cat. No. D9891), clofazimine (CFZ, Cat. No. C8895), dimethyl sulfoxide (DMSO, Cat. No. D2650), Middlebrook Oleic Albumin Dextrose Catalase Growth Supplement (OADC, Cat. No. M0678), Glycerol (Cat. No. G5516), Middlebrook 7H9 Broth Base (Cat. No. M0178), Tween 80 (Polysorbate 80; Cat. No. P5188) and Resazurin sodium salt (Cat. No. R7017) were obtained from Sigma-Aldrich Merck KGaA (Germany), while lyorisenol-3-alpha (LYO-3) and lyorisenol (LYO) were purchased from ChromaDex (USA).

### Bacterial strains and growth conditions

The wild-type WT H37Rv *Mtb* (Pasteur strain) used in this study was generously donated to our team at the Kenya Medical Research Institute (KEMRI) by the *Johnson & Johnson Innovative Medicine*. The BDQ-resistant mutant *Mtb* was spontaneously generated and maintained in glycerol stocks at the *Johnson & Johnson Innovative Medicine*. Cultures were maintained and propagated in Middlebrook 7H9 broth supplemented with 10% OADC, 0.2% glycerol (GLY), and 0.05% Tween80 (Tw80) and on Middlebrook 7H10 agar also supplemented with 10% OADC and 0.2% GLY (7H10_OADC_GLY). Bacteria were subcultured and grown to an early exponential phase prior to experimental use.

### Generation of BDQ-resistant mutants

BDQ-resistant mutants were spontaneously generated as described by (Ismail *et al*., 2018). The WT H37Rv *Mtb* strain was cultured in OADC-supplemented Middlebrook 7H9 broth to an OD₆₀₀ of 1, corresponding to ∼2 × 10⁸ CFU/mL. From this suspension, 1 mL of culture was plated onto Middlebrook 7H10 agar (supplemented with 10% OADC and 0.2% glycerol) containing BDQ at concentrations of 10x, 50x, and 100x MIC_90_ of BDQ, respectively 0.54 µM, 2.7 µM, and 5.4 µM. From this first round of selection, colonies were picked, and clear streaks were made onto 50 x MIC_90_ (2.7 µM) BDQ agar plates to isolate single colonies. After this second round of selection, single colonies were picked to grow in non-selective medium (7H9 supplemented with 10% OADC), for further characterization.

### Resazurin Microtiter Assay (REMA)

The REMA was utilized to determine the MIC of BDQ against WT *Mtb* H37Rv and resistant mutant colonies obtained following BDQ selection as demonstrated by Palomino *et al*., 2002. The working stock of BDQ (12.5 µM) was serially diluted two-fold in supplemented 7H9 broth in sterile 96-well round-bottom microtiter plates. Bacterial cultures were prepared from each resistant colony recovered from BDQ-containing agar plates, along with a wild-type strain as a control. Cultures were adjusted to an OD₆₀₀ of 0.001, and 50 µL of each was added to the wells containing serial drug dilutions, bringing the final assay volume to 100 µL. Plates were incubated at 37 °C for 13 days, after which 20 µL of resazurin was added and incubated for another 24 hours. The MIC was defined as the lowest drug concentration at which no visible growth and no color change from blue to pink were observed. Colonies with MICs four times higher than the WT MIC were classified to be BDQ-resistant according to methods described by (Andries *et al*., 2014) and were selected for downstream genomic analysis.

### Sequencing of mutants for selection of the *Rv0678* mutant

A Hot-Start PCR was performed on BDQ-resistant mutant cultures to amplify the three genes conferring BDQ resistance*: aptE*, Rv0678, and *pepQ* (Yang *et al*., 2020). Primers are listed in Supplementary Table 1. Colonies obtained from BDQ-containing agar plates were picked and cultured in broth to an OD of 1. An aliquot of 1 mL was pelleted and resuspended in 150 µL TE buffer (Invitrogen) as template DNA for the PCR reactions, 2 µL per reaction. The Hot-start PCR reaction began with 15 min at 95°C, followed by a standard PCR program using the Amplitaq Gold 360 DNA Master Mix (Thermo Fisher). These PCR products were purified using the QIAquick kit, verified by gel electrophoresis, and sent to Eurofins for Sanger sequencing with the same primers used for the PCR. Finally, from the selected strains, genomic DNA was extracted using a Quick-DNA Fungal/Bacterial Miniprep kit for whole-genome sequencing to confirm the absence of off-target effects.

### *In vitro* combination studies

A two-dimensional (2D) checkerboard assay was performed to assess the interaction between each EI and BDQ against selected BDQ-resistant *Mtb strain* in a 96-well plate format using resazurin. The fractional inhibitory concentration index (FICI) for each combination was calculated as described by Ramón-García *et al*. 2011.

The FIC for each compound was established using:

FIC_A_ = (MIC of compound A in the presence of compound B) / (MIC of compound A alone), where FIC_A_ represents the fractional inhibitory concentration of compound A. Similarly, the FIC for compound B (FIC_B_) was calculated. The overall FICI was the sum of FIC_A_ and FIC_B_. The interaction was classified based on the FICI values: synergy was described as FICI ≤ 0.5, antagonism as FICI > 4.0, and no interaction for FICI values between 0.5 and 4.0.

### Minimum bactericidal concentration (MBC) assay

Wells indicative of synergism from the best BDQ-EI combination against the BDQ-resistant *Mtb* strain were selected for MBC determination. 100 µL aliquots of the synergy well contents were aseptically aspirated, diluted 10-fold in sterile phosphate buffer and plated onto drug-free 7H10_OADC_GLY agar infused with activated charcoal. Colony enumeration was made after incubation of the plates at 37^0^C for 21 days. The MBC was defined as the lowest concentration of the BDQ-EI combination at which no colony growth was observed, indicating ≥99.9% bacterial killing. This step was critical in confirming the sterilizing potential of the BDQ-EI combination following the inhibition data obtained from the checkerboard assay.

## Results

### *In vitro* selection of Rv0678 mutants under BDQ pressure

An attempt to generate spontaneous BDQ-resistant strains of *Mtb* was successful. The MIC of BDQ against the WT H37Rv *Mtb* was determined to be 0.098µM as shown in Table 2. The starting inoculum of WT *Mtb* H37Rv had been standardized to ∼1.0 x 10⁸ CFU/mL to allow accurate determination of resistance frequency. Following exposure, bacterial growth was detected on agar plates containing elevated BDQ concentrations, indicating the presence of spontaneous resistant mutants that survived BDQ pressure. Plates containing 10x the MIC of BDQ showed confluent growth with over 100 colonies; therefore, an accurate calculation of the resistance frequency could not be performed. At 50x MIC BDQ, an average of 18 resistant colonies were identified, yielding an estimated resistance frequency of 1.8 x 10⁻⁷. Across replicate plates containing 100xMIC BDQ, an average of 7 colonies per plate were recovered, corresponding to a resistance frequency of ∼7.0 x 10⁻⁸ (Table 1). From all BDQ-containing plates (50x and 100x MIC conditions), a total of 18 colonies were randomly selected as potential resistant mutants for MIC evaluation to assess their BDQ resistance profiles.

**Table 1.**
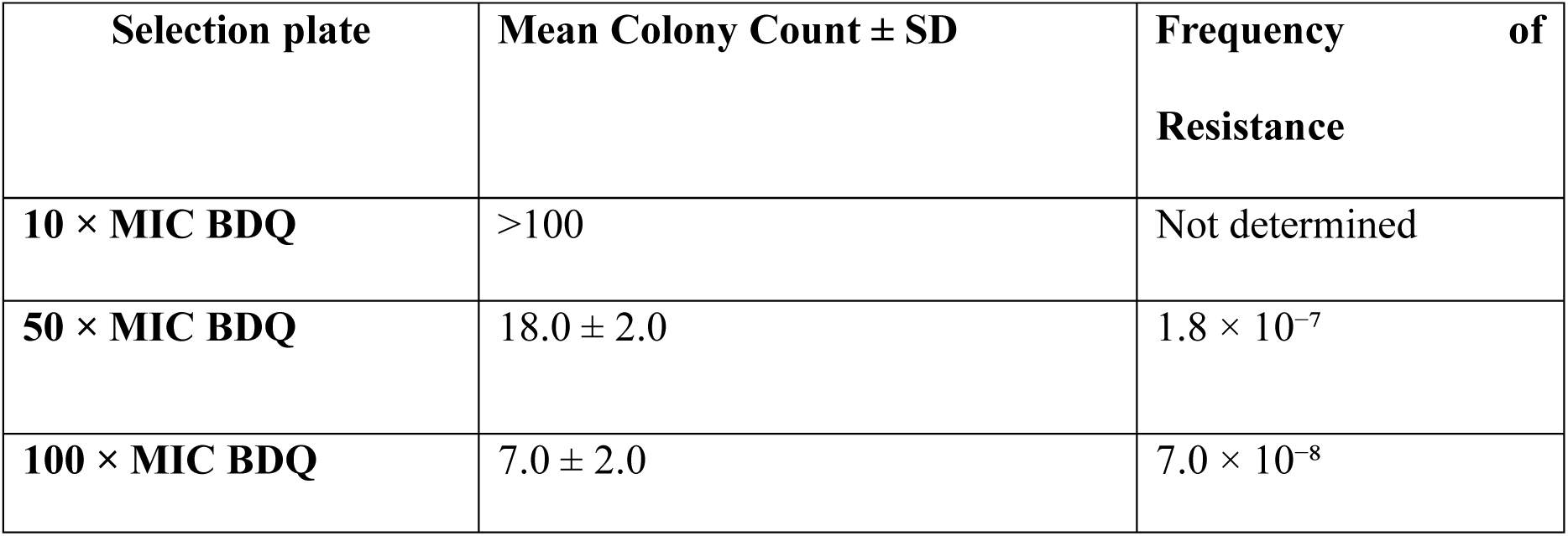
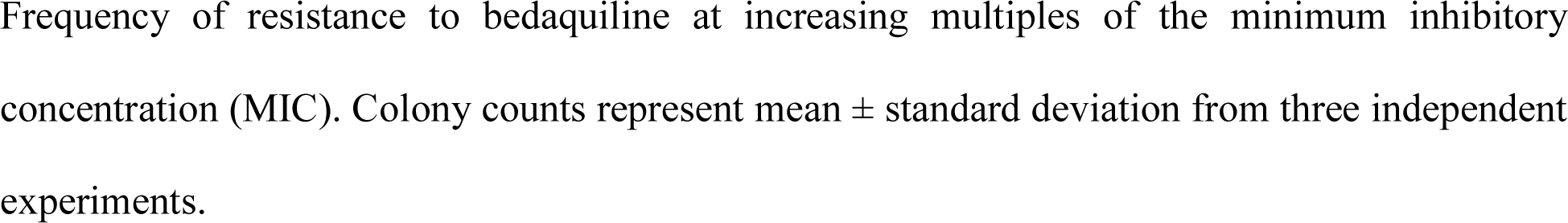
Colony counts and resistance frequencies of spontaneous BDQ-resistant *Mtb* mutants at different BDQ concentrations.

| Selection plate | Mean Colony Count ± SD | Frequency of Resistance |
| --- | --- | --- |
| 10 × MIC BDQ | >100 | Not determined |
| 50 × MIC BDQ | 18.0 ± 2.0 | 1.8 $\times 10^{-7}$ |
| 100 × MIC BDQ | 7.0 ± 2.0 | 7.0 $\times 10^{-8}$ |

Frequency of resistance to bedaquiline at increasing multiples of the minimum inhibitory concentration (MIC). Colony counts represent mean ± standard deviation from three independent experiments.

**Table 2:** Descriptive Statistics for the MIC of BDQ against spontaneously generated BDQ-resistant *Mtb* strains.

| Group | n | Mean MIC<br>(μM) | Median (μM) | SEM | Range (μM) |
| --- | --- | --- | --- | --- | --- |
| Drug free | 3 | 0.098 | 0.098 | 0.000 | 0.098 – 0.098 |
| BDQ 10x | 3 | 0.131 | 0.098 | 0.033 | 0.098 – 0.196 |
| BDQ 50x | 27 | 0.725 | 0.392 | 0.120 | 0.196 – 3.136 |
| BDQ 100x | 12 | 1.027 | 0.784 | 0.237 | 0.196 – 3.136 |

Summary of MIC values for BDQ against mutant strains (n=14) and the WT H37Rv reference strain, presented as mean ± standard deviation, median, and range.

### Validation of BDQ-resistant phenotype in spontaneously generated mutants

Of the 18 emerging colonies, only 14 had their MIC against BDQ evaluated, and 4 were eliminated because they could no longer grow. The WT strain showed uniform susceptibility, with MICs of 0.098 μM across all repeats. The single mutant (1) selected at 10x (0.98 μM) had MICs very close to the WT (0.098 μM - 0.196 μM). Mutants selected from 50x MIC BDQ had higher MICs ranging from 0.392 μM to 0.784 μM, which were 4- to 8-fold higher than WT. Some selected colonies, such as mutants 11 and 13 from agar plates with a BDQ concentration of 50x (4.9 µM), exhibited an MIC of 3.136 µM, which was 32-fold higher than that of the WT. Similarly, mutants from the 100x MIC BDQ selection had elevated MICs ranging from 0.392 μM to 3.136 μM, with several exhibiting strong resistance, as shown in Table 2 and Figure 1.

**Figure 1:**
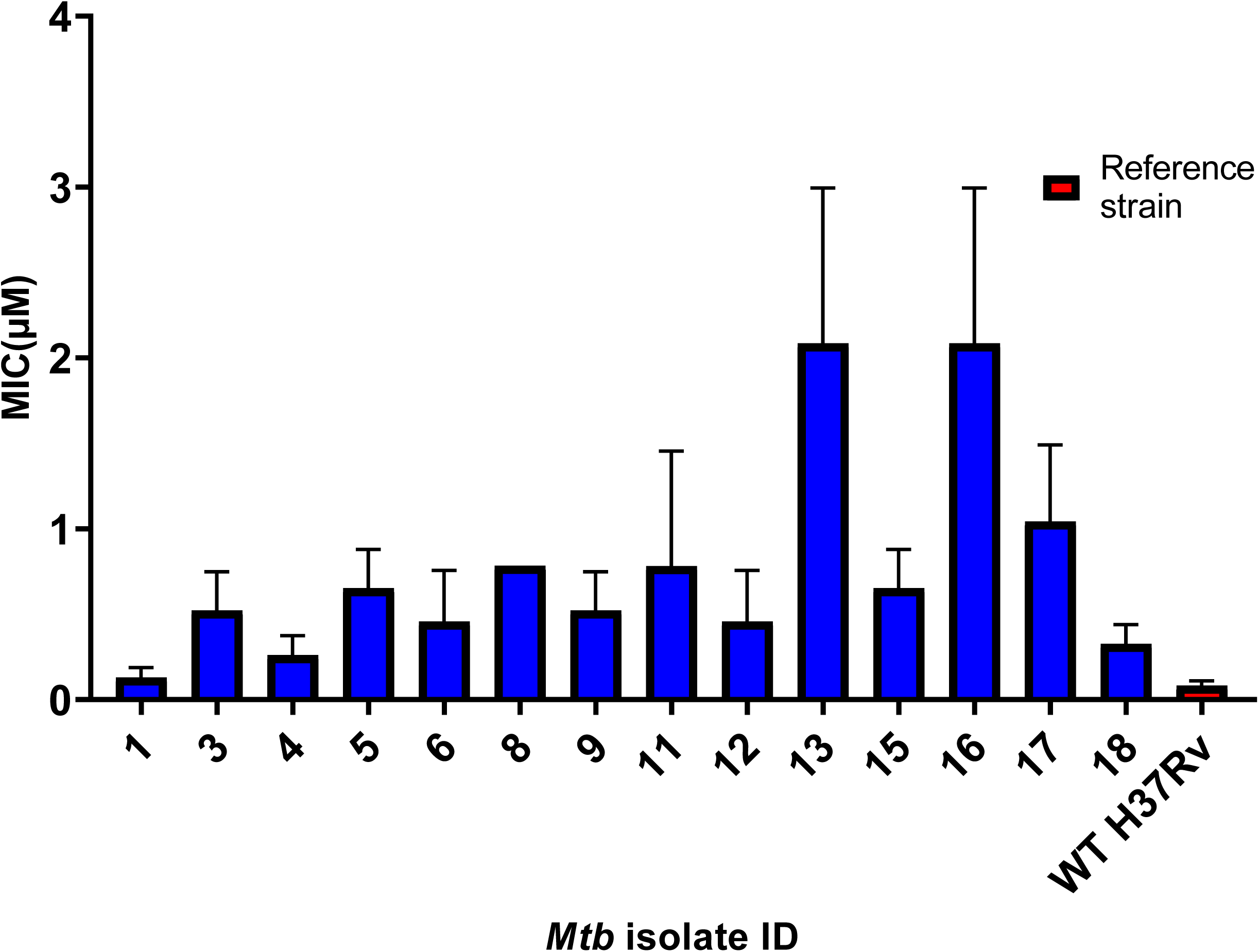
The MIC of resistant selected colonies (in blue) compared with the WT H37Rv strain (in red). The figure visually displays the MIC values for each selected colony, along with the reference strain used in the study. WT strain grown on BDQ-free agar served as a control and reference strain. The MIC values BDQ-resistant colonies show an increase ranging from 2 to 32 times when compared to the MIC of the WT H37Rv strain.

### Validation of the BDQ-resistant genotype in spontaneously generated mutants

All 14 BDQ-resistant *Mtb* colonies (1, 3, 4, 5, 6, 8, 9, 11, 12, 13, 15, 16, 17, and 18), along with the wild-type H37Rv strain, were successfully amplified. The expected amplicon sizes were consistently observed: approximately 649 base pairs (bp) for *Rv0678*, 1334 bp for *pepQ*, and 522 bp for *AtpE*. Each colony produced a single, well-defined band per gene, with no evidence of smearing or nonspecific amplification (Figure 2). The banding patterns were uniform in appearance, and the amplification products were of comparable clarity and intensity across all samples, suggesting robustness of the methodology. Importantly, the WT H37Rv strain exhibited identical amplicon sizes, confirming PCR specificity and providing a baseline for evaluating gene integrity in resistant strains. The presence of all three gene bands across the different selected colonies, as seen on panels a, b, c, and d (Figure 2), regardless of the BDQ concentration used during selection (10x, 50x, or 100x), suggested intact gene regions suitable for downstream sequence analysis. The sequencing revealed a mixture of point mutations and frameshift alterations (Table 3). In the *Rv0678* gene, several point mutations were detected. For instance, mutant 1 showed a thymine-to-cytosine substitution at the first codon, leading to a methionine-to-alanine amino acid change. Selected colonies 4 and 12 harbored a guanine-to-thymine mutation at codon 66, potentially resulting in either a glycine-to-valine or methionine substitution. Colony 11 carried a cytosine-to-thymine mutation at codon 94, which potentially led to an arginine-to-tryptophan substitution. In addition to these point mutations, frameshift mutations were observed in selected colonies 5 and 13. These were caused by a single-base deletion and a 17-base pair insertion, respectively. Both mutations introduced premature stop codons at amino acid position 32, highly likely to disrupt the function of the *Rv0678* gene product. Another frameshift in colony 6 was caused by a cytosine insertion at codon 156, extending the open reading frame past the natural stop codon and potentially altering protein function. In the *atpE* gene, a guanine-to-thymine mutation at codon 61 was consistently observed in five resistant-selected colonies (3, 15, 16, 17, and 18), potentially leading to a glutamic acid-to-aspartic acid substitution. For the *pepQ* gene, a cytosine-to-guanine mutation at codon 66 was detected in selected colonies 8 and 9, resulting in an isoleucine-to-methionine substitution. However, due to poor sequencing quality, results from selected colonies 1 and 13 could not be reliably interpreted for *pepQ*.

**Figure 2:**
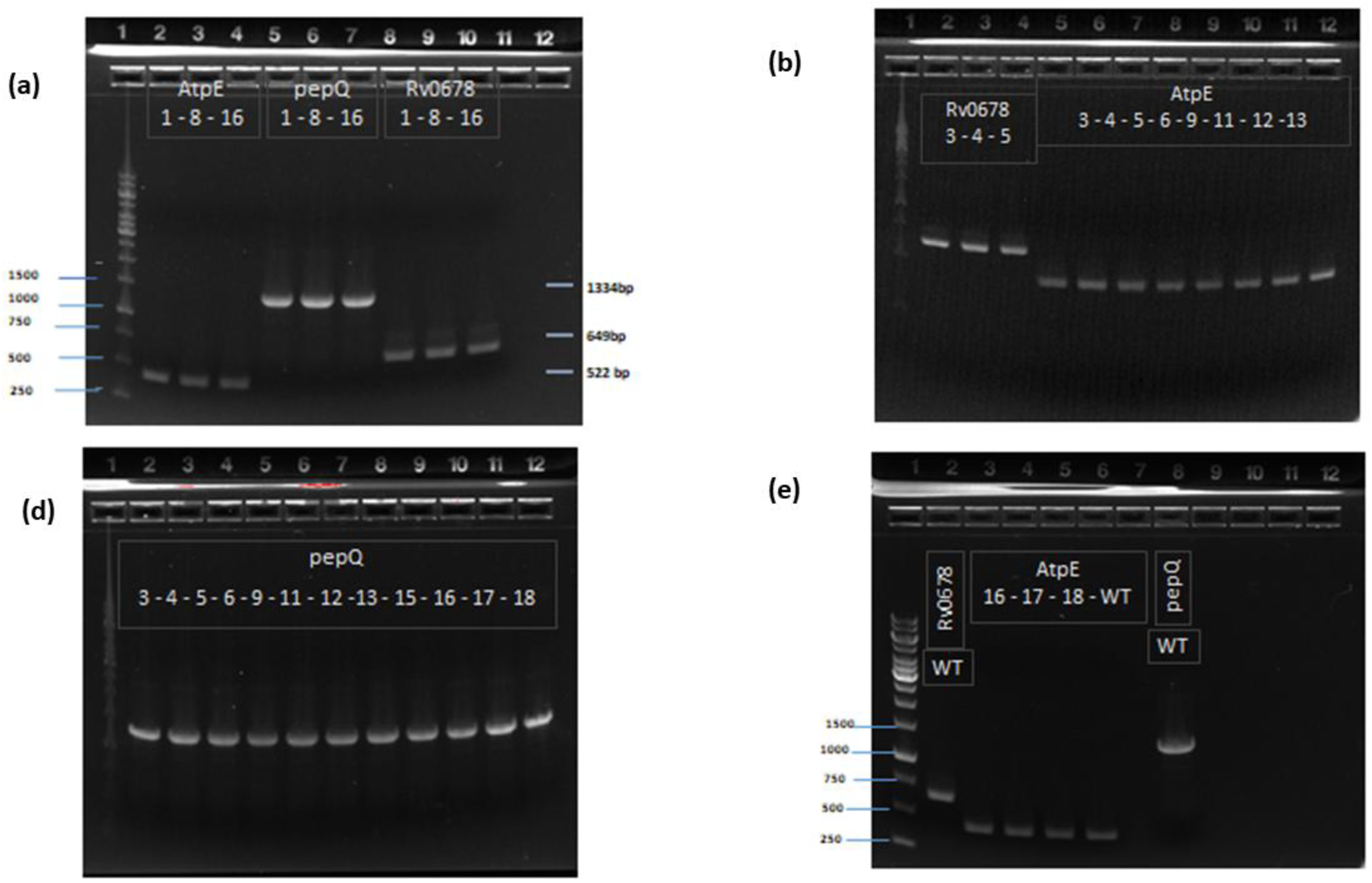
Agarose gel electrophoresis showing successful PCR amplification of the targets of interest; (**a**) PCR amplification of *AtpE* (522 bp), *pepQ* (1134 bp), and *Rv0678* (649 bp) from DNA isolated from colonies 1, 8, and 16; (**b**) Amplification of *Rv0678* and *AtpE* in colonies (3-13); **(c)** Amplification of *pepQ* region across colonies 3-18; **(d)** Amplification of *Rv0678, AtpE*, and *pepQ* in WT H37Rv and resistant selected colonies 16–18; WT denotes the wild-type control.

**Table 3:**
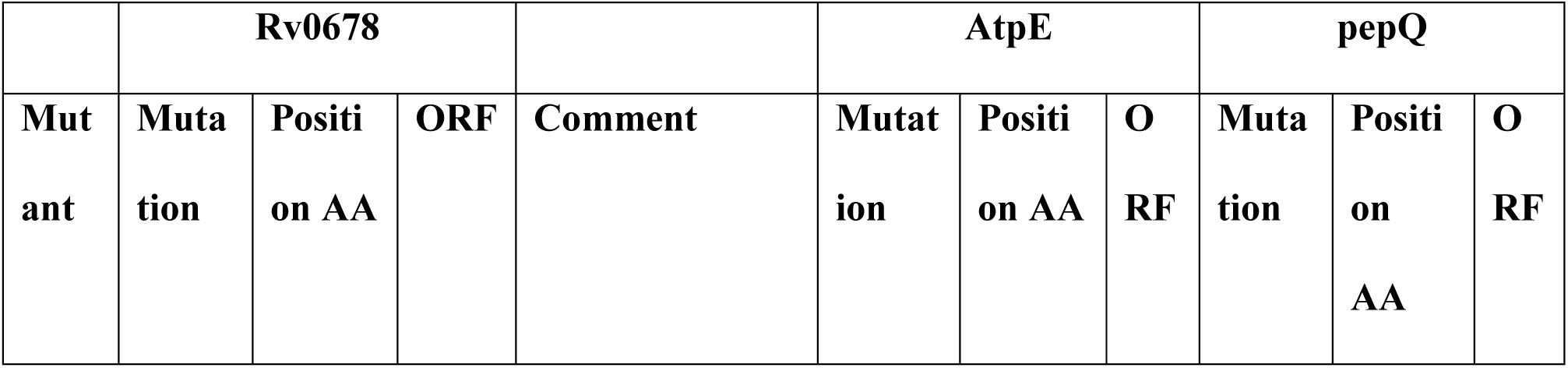
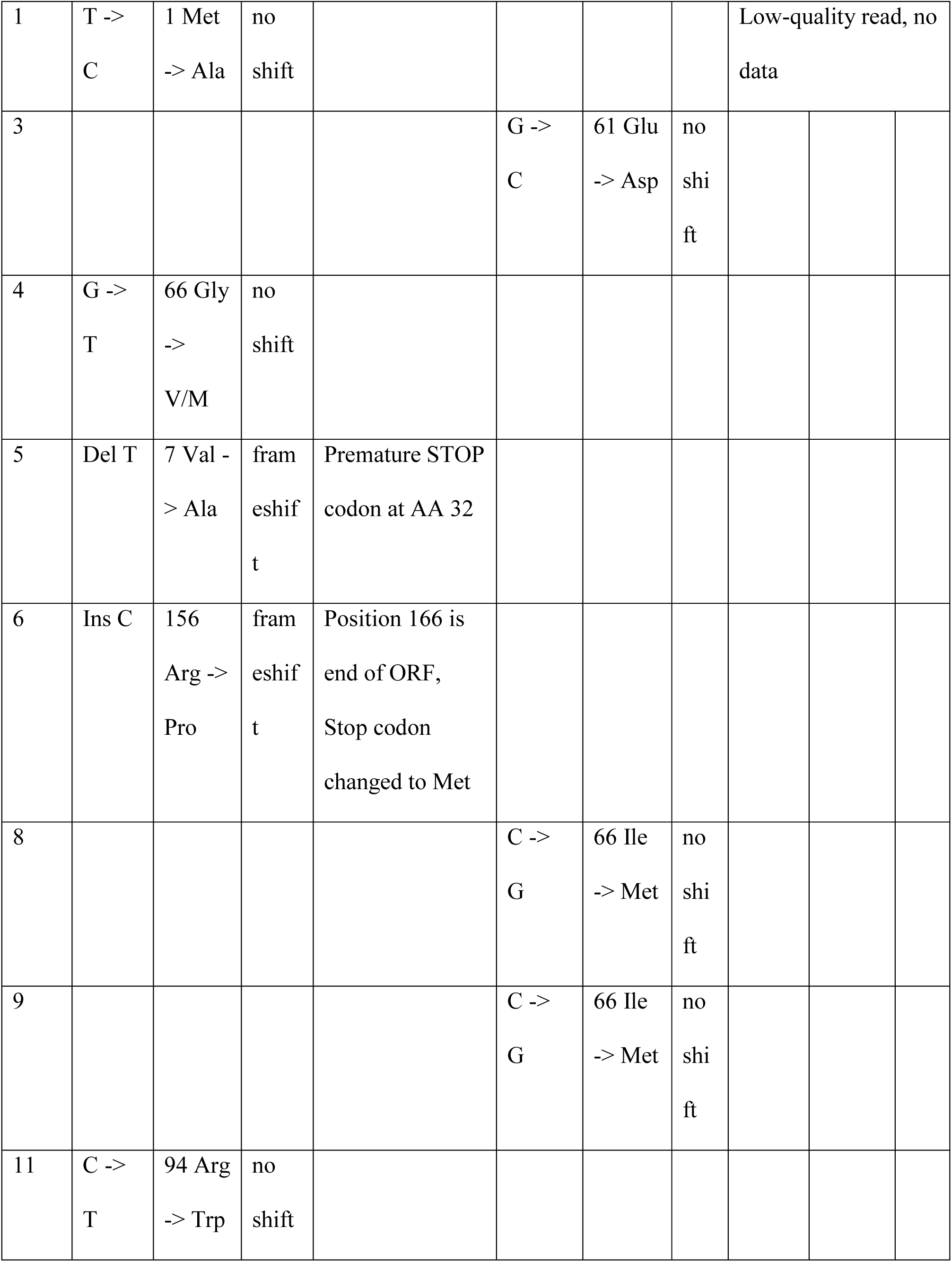
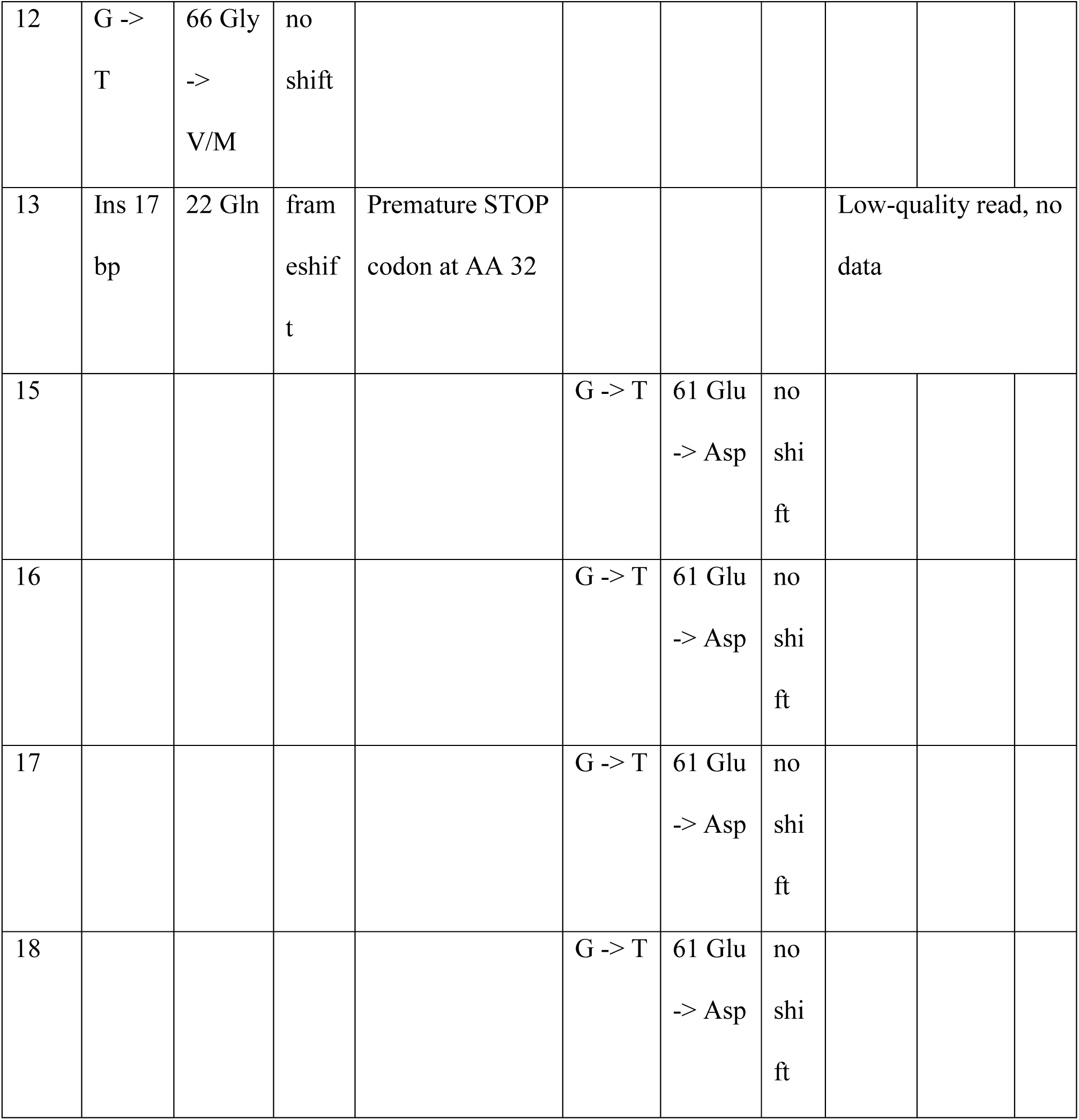
Summary of nucleotide and amino acid changes detected in *Rv0678*, *atpE*, and *pepQ* genes. **Table 3:** Summary of nucleotide and amino acid changes detected in *Rv0678, atpE*, and *pepQ* genes among 14 spontaneously generated *Mtb* selected colonies; T-Thymine; C-cytosine; G505 Guanine; AA-amino acid; ORF- open reading frame; Glu-glutamic acid; Asp-aspartic acid; Gln506 glutamine; Arg-Arginine; Gly-glycine; Trp-tryptophan; Met-methionin; Ala-alanine; Val-valine; Pro-proloine; bp-base pair; Ins-insertion; Del-deletion; V/M-valine/methionine; Ile-isoleucine

|  | Rv0678 |  |  |  | AtpE |  |  | pepQ |  |  |
| --- | --- | --- | --- | --- | --- | --- | --- | --- | --- | --- |
| Mutant | Mutation | Position AA | ORF | Comment | Mutation | Position AA | ORF | Mutation | Position AA | ORF |
| 1 | T -><br>C | 1 Met<br>-> Ala | no<br>shift |  |  |  |  | Low-quality read, no data |  |  |
| 3 |  |  |  |  | G -><br>C | 61 Glu<br>-> Asp | no<br>shi<br>ft |  |  |  |
| 4 | G -><br>T | 66 Gly<br>-><br>V/M | no<br>shift |  |  |  |  |  |  |  |
| 5 | Del T | 7 Val -<br>> Ala | fram<br>eshif<br>t | Premature STOP<br>codon at AA 32 |  |  |  |  |  |  |
| 6 | Ins C | 156<br>Arg -><br>Pro | fram<br>eshif<br>t | Position 166 is<br>end of ORF,<br>Stop codon<br>changed to Met |  |  |  |  |  |  |
| 8 |  |  |  |  | C -><br>G | 66 Ile<br>-> Met | no<br>shi<br>ft |  |  |  |
| 9 |  |  |  |  | C -><br>G | 66 Ile<br>-> Met | no<br>shi<br>ft |  |  |  |
| 11 | C -><br>T | 94 Arg<br>-> Trp | no<br>shift |  |  |  |  |  |  |  |
| 12 | G -><br>T | 66 Gly<br>-><br>V/M | no<br>shift |  |  |  |  |  |  |  |
| 13 | Ins 17<br>bp | 22 Gln | fram<br>eshif<br>t | Premature STOP<br>codon at AA 32 |  |  |  | Low-quality read, no<br>data |  |  |
| 15 |  |  |  |  | G -> T | 61 Glu<br>-> Asp | no<br>shi<br>ft |  |  |  |
| 16 |  |  |  |  | G -> T | 61 Glu<br>-> Asp | no<br>shi<br>ft |  |  |  |
| 17 |  |  |  |  | G -> T | 61 Glu<br>-> Asp | no<br>shi<br>ft |  |  |  |
| 18 |  |  |  |  | G -> T | 61 Glu<br>-> Asp | no<br>shi<br>ft |  |  |  |

### Rescue effect of efflux inhibitors in combination with BDQ in BDQ-resistant mutant

Colony 13 was selected for this experiment as it had a frameshift in the Rv0678 gene, resulting in a premature stop codon, which could possibly be causing an overexpression of the MmpS5-MmpL5 pump, causing resistance (Ma *et al*., 2020). This is evidenced by the high MIC values of BDQ as shown in Table 2. The MIC of BDQ against the WT H37RV strain was 0.098 µM, whereas for the BDQ-resistant mutant 13, a 16-fold increase in the MIC (1.565 µM) was observed, indicating decreased drug sensitivity. When combined with BER, the activity of BDQ against the resistant strain was reduced from 1.563 µM to 0.098 µM, representing a 16-fold improvement and a chemical rescue of the phenotype. Notably, BER alone exhibited activity, with an MIC₉₉ of 201.67 µM, against the mutant. The Fractional Inhibitory Concentration Index (FICI) for this combination was 0.188. Again, a combination of RES with BDQ against BDQ mutant 13 lowered the MIC₉₉ of the variant from 1.563 µM to 0.195µM, an 8-fold change, and with FICI of 0.386, suggesting a synergistic effect. The combination of BDQ with PIP resulted in a 16-fold change, lowering the BDQ mutant MIC₉₉ from 1.563 µM to 0.098 µM. The MIC₉₉ of PIP tested singly was 0.077 µM, and the FICI for this pair was 0.557, just above the threshold for synergy, indicating an additive effect. Another ideal synergistic result was observed when BDQ was combined with LYO, which reduced the MIC from 1.563 µM to 0.002 µM, a 64-fold reduction, as shown in Table 4 and Figure 3. Lastly, the BDQ and LYO-3 combination also achieved a 16-fold reduction in the MIC₉₉ of the BDQ mutant, from 1.563 µM to 0.39 µM.

**Table 4:**
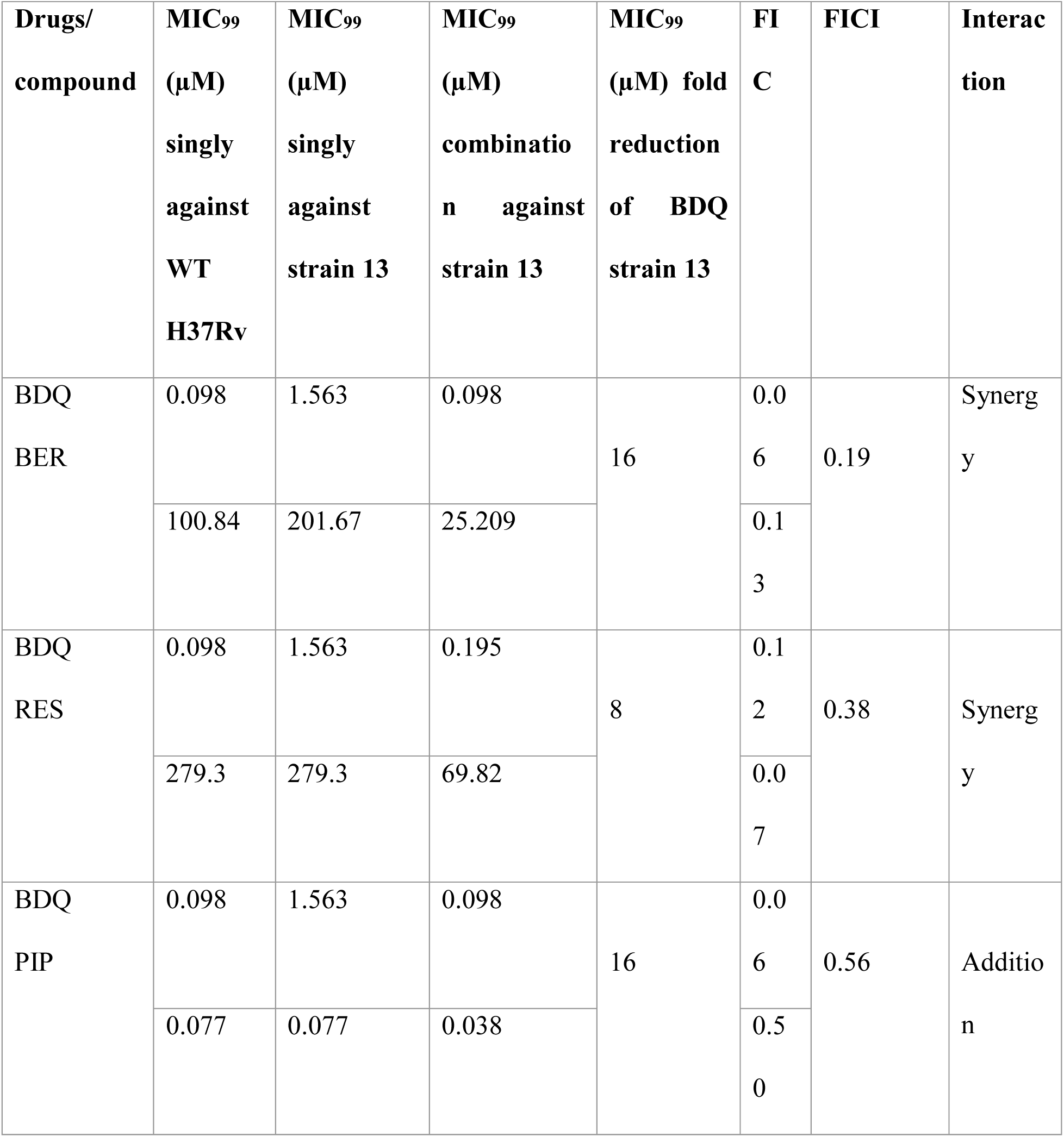
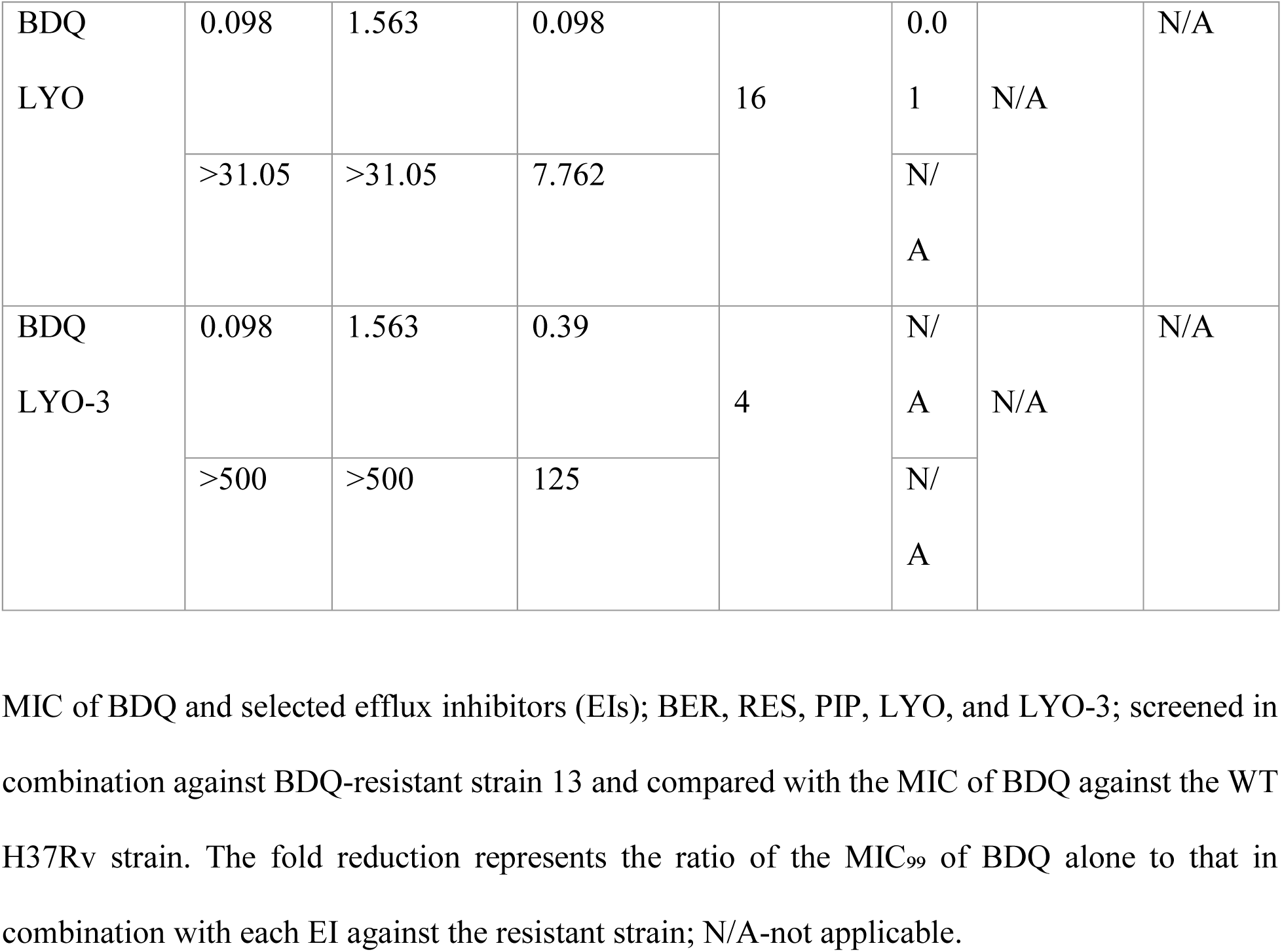
MIC of BDQ when combined with efflux inhibitors (EIs) against BDQ-resistant mutant strain 13.

**Figure 3.**
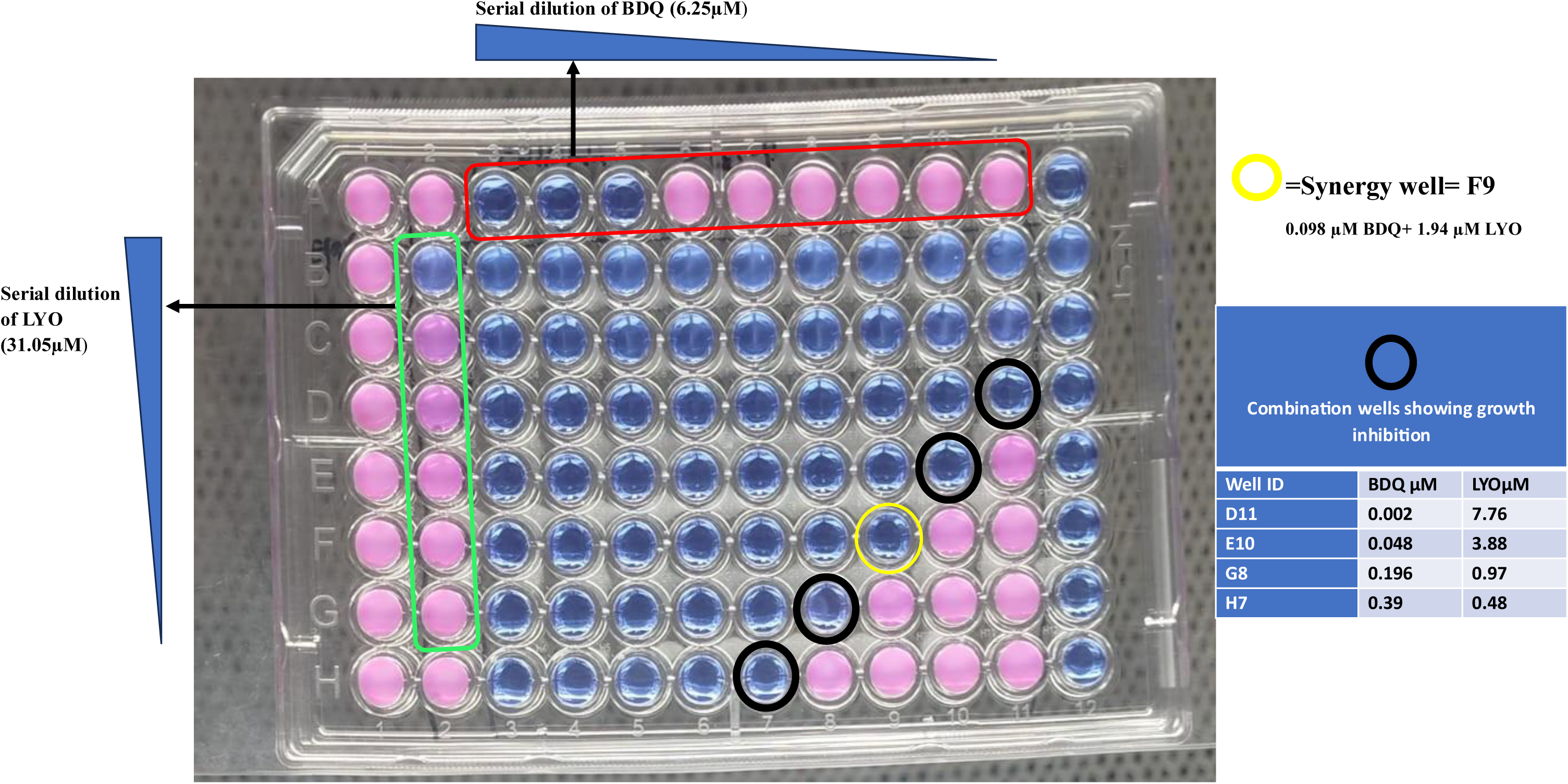
Checkerboard Assay of BDQ and LYO against Resistant *Mtb* Mutant 13. A 96-well microtiter plate illustrating the interaction between BDQ and LYO in a two-dimensional serial dilution format. BDQ was serially diluted horizontally across the columns (starting at 6.25 µM; top row highlighted in red), while LYO was serially diluted vertically down the rows (starting at 31.05 µM; left column highlighted in green). Each well represents a unique combination of BDQ and LYO concentrations. Pink wells indicate viable *Mtb,* and blue wells show inhibited growth. Column 1 was the growth control (no drug, pink wells), and column 2 was the sterility control (no bacteria, blue wells). The yellow-circled well (F9) highlights the synergistic interaction (Ramón-García et al., 2011), when BDQ and LYO are combined.

### Minimum Bactericidal Concentration (MBC) of BDQ–LYO Synergy

Having a comparable fold reduction to EIs that have previously been reported (Jin *et al*., 2011; Morita *et al*., 2016; Shaheen *et al*., 2019). The LYO+BDQ combo was selected for the MBC assay. Contents from all drug-combination wells that showed growth inhibition, as highlighted in Figure 3, were aspirated. Dilutions were made, and plating was done as shown in panels (a), (b), (c), and (d) in Figure 4. Emerging CFUs were counted after 21 days of incubation at 37°C, and the results are summarized in Table 5. The untreated inoculum averaged 2.05 x 10⁸ CFU/mL, establishing the baseline bacterial load. In the growth control (A2), where no drug was applied, bacterial proliferation was evident, with counts increasing to 1.12 x 10¹⁰ CFU/mL. This confirmed both the culture’s viability and the suitability of the assay conditions. At the lowest tested concentration (D11: 0.002µM BDQ + 7.76µM LYO), bacterial counts remained comparable to the inoculum (2.09 x 10⁸ CFU/mL), corresponding to a 1-log_10_ reduction. Increasing the BDQ concentration to 0.048µM (E10), in combination with 3.88µM LYO, resulted in a further decline to 1.30 x 10⁶ CFU/mL, representing a 2-log_10_ reduction. A further transition toward CFU reduction was observed at F9, the synergy well (0.098 µM BDQ, 1.94 µM LYO), where bacterial counts dropped to 4.24 × 10⁵ CFU/mL, corresponding to a 3-log_10_ reduction. This trend intensified at G8 (0.196 µM BDQ, 0.97 µM LYO), yielding 2.58 x 10⁴ CFU/mL and a 4-log_10_ reduction, indicating strong killing activity. The most pronounced effect was observed at the highest tested concentration (H7: 0.39 µM BDQ, 0.48 µM LYO), where bacterial counts collapsed to 1.12 x 10³ CFU/mL, corresponding to a 5-log_10_ reduction.

**Figure 4:**
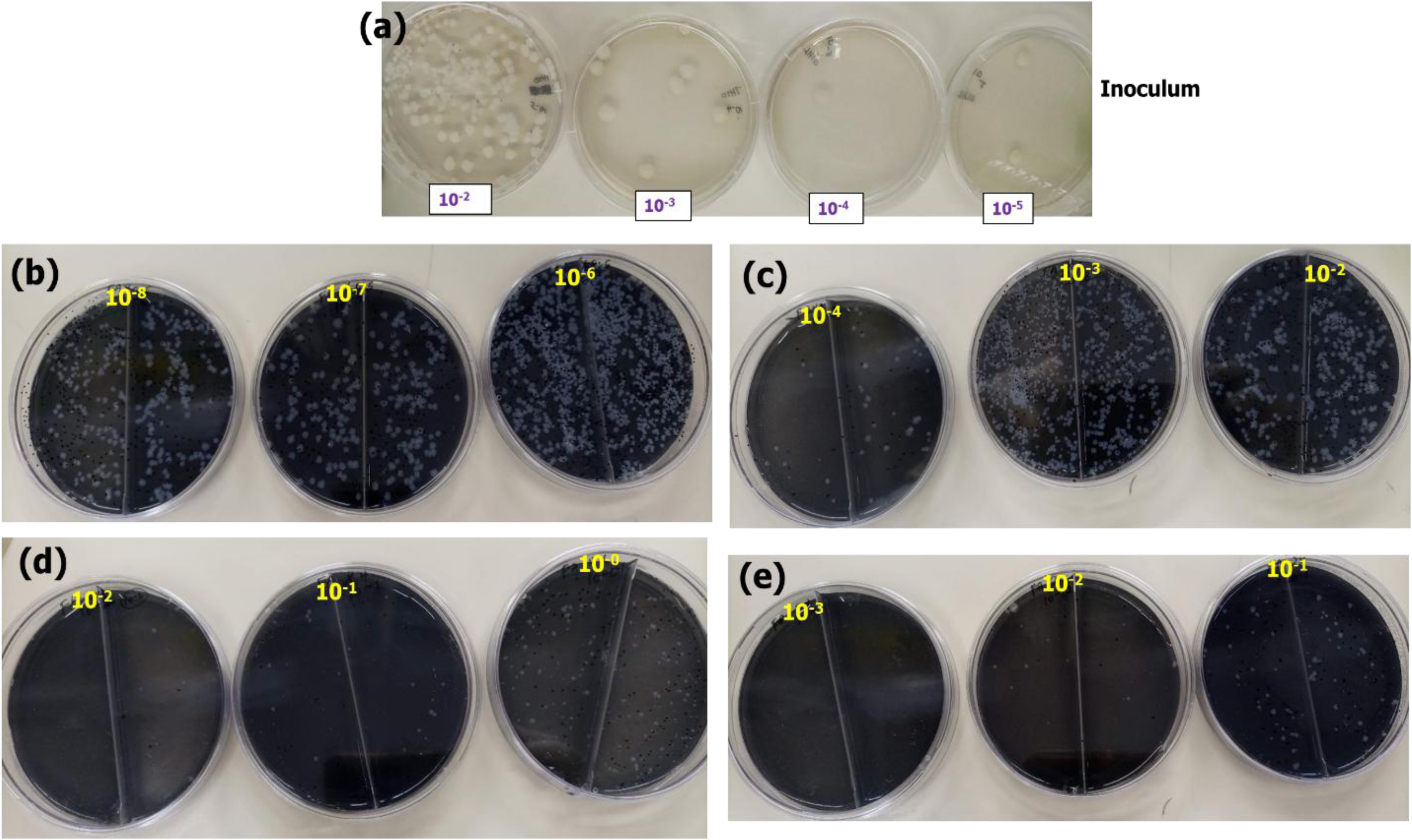
CFU of *Mtb* plated from BDQ+LYO wells that showed growth inhibition following the checkerboard assay in Figure 3. **(a)** Well contents from untreated control (well A2); **(b)**, **(c)**, **(d),** and **(e)**-well contents from wells D11, F9, G8, and H7, respectively.

**Figure 5:**
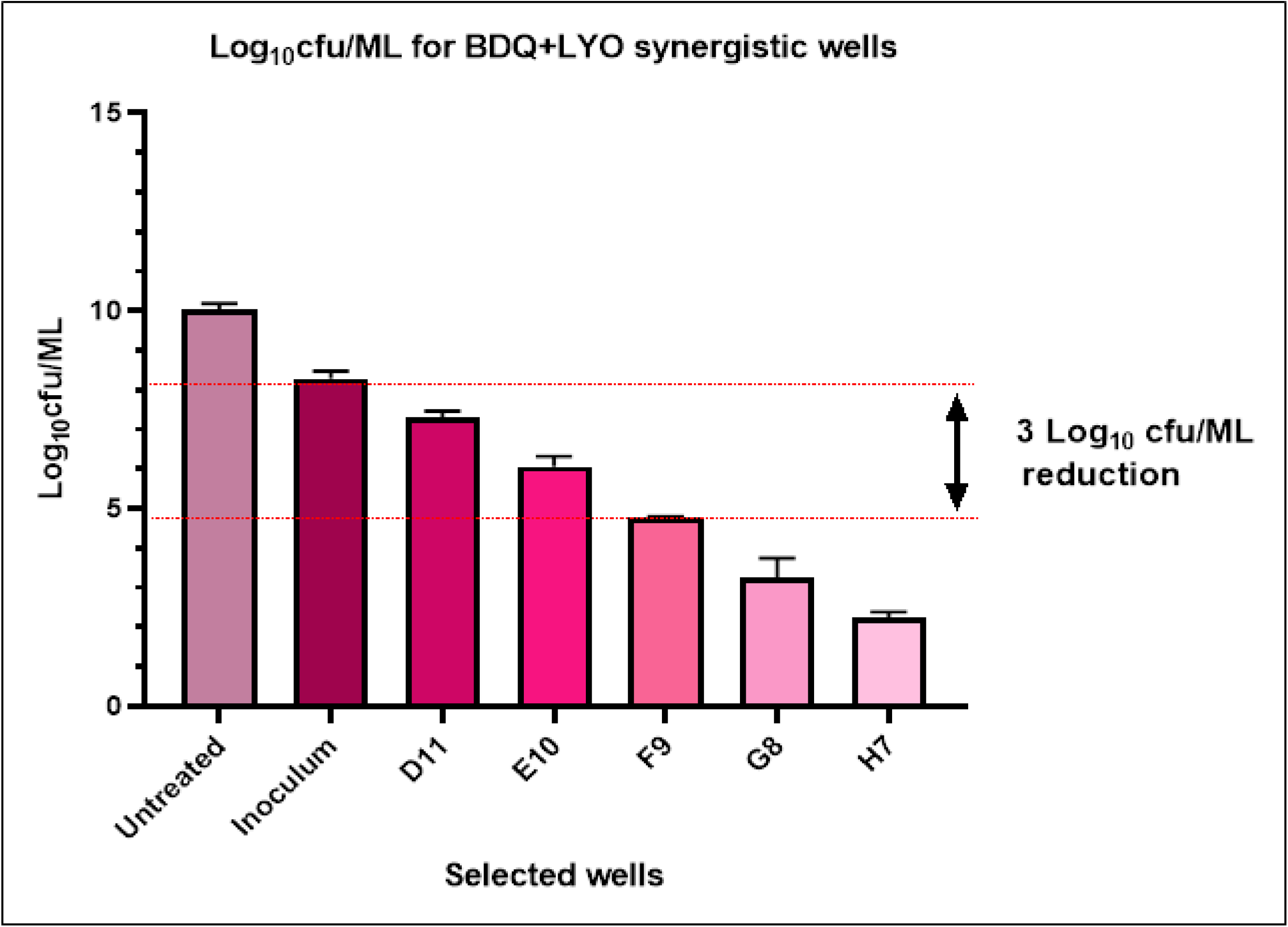
Log_10_ CFU/mL for selected BDQ+LYO wells from the two-dimensional checkerboard assay in Figure 3. The bar graph shows log₁₀ CFU/mL for the untreated control, inoculum, and wells D11 to H7 after treatment with BDQ plus lyoniresinol (LYO). D11/E10 exhibits bacteriostatic reductions, while F9, G8 and H7 show progressive bactericidal activity, with G8 ≥3-log_10_ and H7 ≥5-log_10_ decreases. Dashed red lines mark log_10_-reduction thresholds.

**Figure 6:**
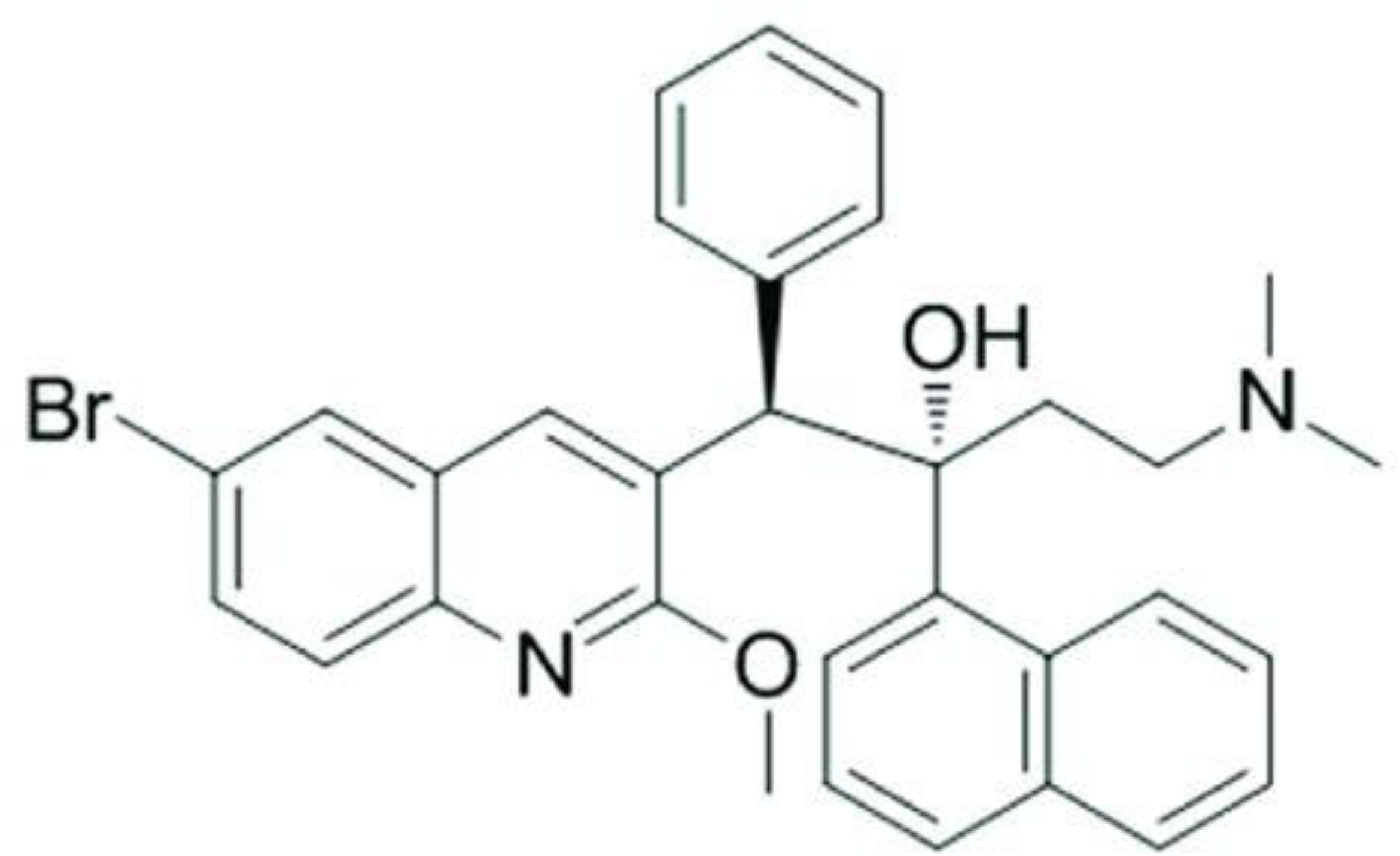
Structure of Bedaquiline

**Figure 7:**
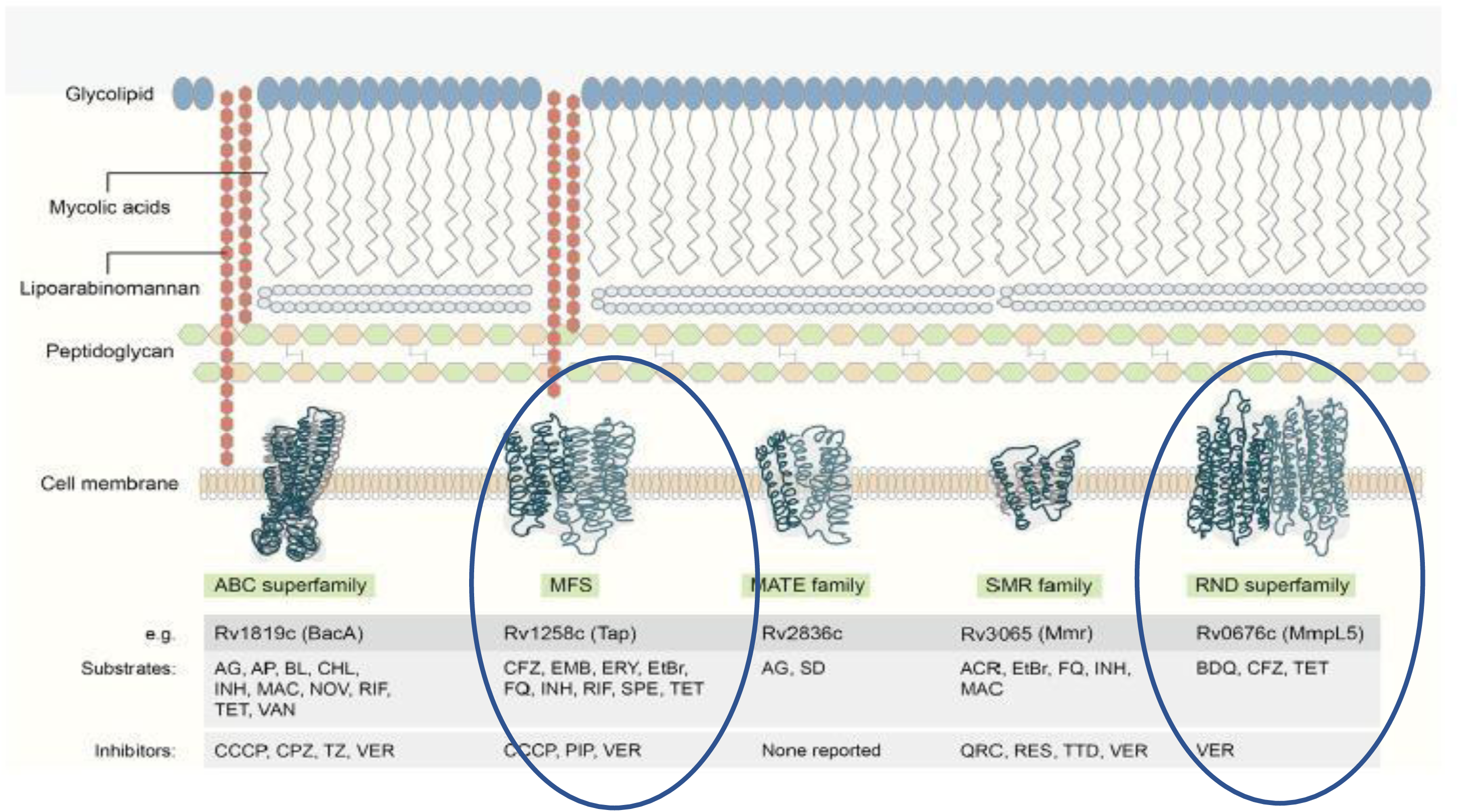
A demonstration of *Mtb* Efflux Pumps by Laws *et al.,* 2022

**Table 5:**
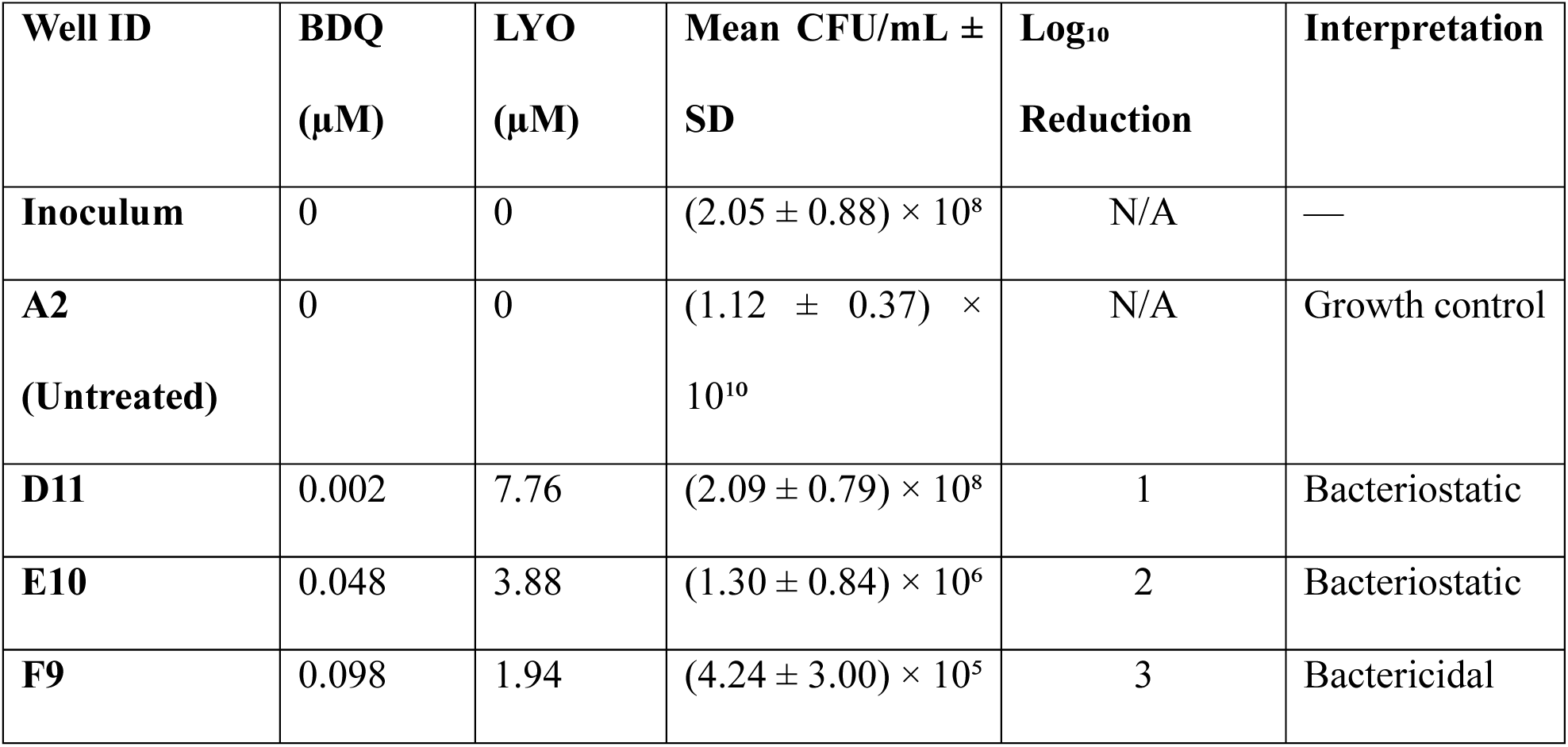
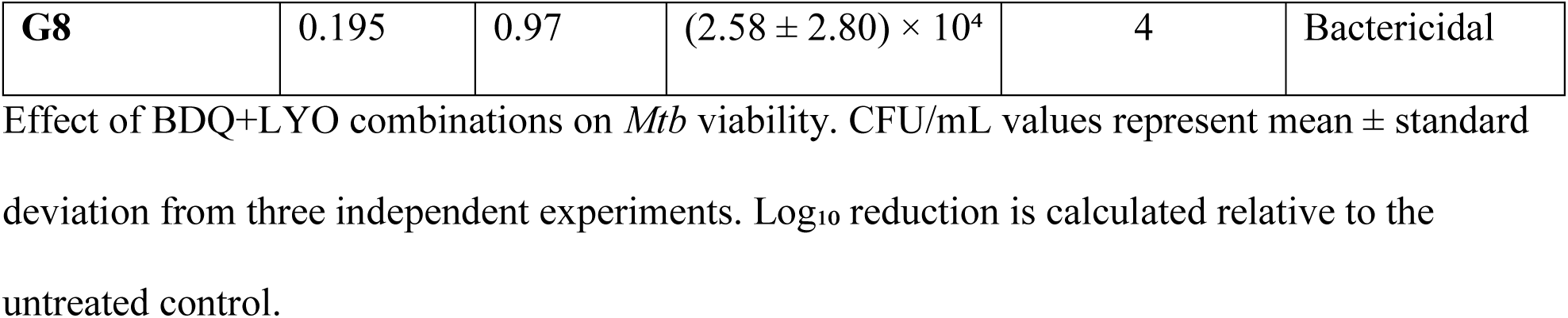
MBC of selected synergistic wells of BDQ and LYO combinations against BDQ-resistant mutant 13.

## Discussion

Bedaquiline (BDQ) is the pillar of current anti-TB chemotherapy for drug-resistant tuberculosis (Goletti *et al.,* 2025). However, the long-term clinical utilization of this drug is threatened by the emergence and spread of *Mtb Rv0678* mutations that upregulate the MmpS5-MmpL5 pump. These alterations interfere with the repressor protein’s normal activity, rendering it incapable of binding to DNA and blocking transcription of EP genes. As a result, efflux pump expression is upregulated, enabling bacteria to expel antimicrobial agents more effectively (Xu *et al*., 2023). This study aimed to generate BDQ resistance in *Mtb* and use the resulting mutant as a model for *in vitro* evaluation of whether selected natural product-derived EIs can functionally rescue BDQ resistance. By generating resistant strains under BDQ pressure, identifying genetic mutations associated with resistance, and evaluating drug interactions with EIs, this investigation aimed to assess the capacity of efflux inhibitors to restore susceptibility to BDQ in *Mtb-*resistant strains. The emergence of *Mtb* colonies on medium containing high BDQ concentrations suggested the successful generation of spontaneous mutants that survived these high concentrations, likely due to mutations. This finding aligns with observations by (Ismail *et al*., 2018), who reported that *Mtb* can develop mutations that enable it to survive and grow in the presence of high drug concentrations. These adaptations may explain the growth observed in drug-containing media despite the high levels used.

Resistance frequency declined with increasing BDQ levels, although colonies still appeared at 9.8 µM, underscoring *Mtb*’s robust ability to adapt under drug pressure despite its inherently slow growth rate (Xu *et al*., 2023). These results suggest that exposure to high drug doses may be insufficient to kill all bacteria and can select for and promote the growth of resistant mutants. As such, higher bacterial numbers may emerge, increasing the probability that resistance evolves and persists (Edwards & Field, 2022). Mutant colonies selected at lower BDQ levels showed MICs comparable to the wild type, while those obtained from 50x and 100x BDQ plates exhibited MICs up to 32-fold higher. This trend aligns with findings by Andries *et al*. (2014), who demonstrated a model of stepwise resistance acquisition and indicated that even minor fluctuations in environmental drug concentration can promote the selection of progressively less-susceptible mutants. In this study, the three key genes previously reported (Islam *et al*., 2024) to contribute to BDQ resistance were found to harbor mutations in all 14 selected mutant colonies, making them suitable for drug testing/screening.

The frameshift mutation observed in mutant colonies 5 and 13 introduced a premature stop codon early in the Rv0678 open reading frame. This alteration not only disrupted the reading frame, leading to a completely different downstream amino acid sequence, but also caused early translation termination, resulting in a truncated, nonfunctional protein. Even if partial resemblance to the wild-type Rv0678 were retained, the premature stop codon ensures loss of repressor integrity. Consequently, these mutations strongly suggest complete abrogation of Rv0678 repressor function, leading to de-repression and overexpression of the MmpL5-MmpS5 efflux pump. This mechanistic explanation is consistent with previous reports showing that truncating or modifying mutations in Rv0678 abolish repressor activity, thereby enhancing BDQ efflux and reducing intracellular drug concentrations (Limberis *et al*., 2023). The elevated MIC values observed in these mutants are therefore likely attributable to disruption of regulatory control and subsequent hyperactivity of the efflux system.

Additionally, point mutations in this gene, such as Arg94Trp in colony 11, have previously been reported to compromise the DNA-binding and structural integrity of the protein, further promoting excessive efflux pump expression in clinical BDQ-resistant strains (Rashid *et al*., 2025). The *atpE* gene encodes the target of BDQ, the c subunit of ATP synthase. Mutations in the gene reduce drug binding and confer high-level resistance (S *et al*., 2020). Sequencing revealed a recurring Glu61Asp mutation in *atpE* appearing in several mutant colonies. Although conservative, this mutation may have subtly altered the drug’s binding site, reducing its effectiveness. The *pepQ* gene, which likely encodes an aminopeptidase involved in protein metabolism, has been linked to low-level resistance; however, its precise role remains unclear (Almeida *et al*., 2016). Mutations in *pepQ* were rare, though an Ile66Met change was detected in mutant colonies 8 and 9. Collectively, these outcomes support the assumption that the loss-of-function mutations in the Rv0678 gene is vital for efflux-mediated BDQ resistance, and that alterations in *atpE* and *pepQ* may confer additional resistance or modulate efflux activity. As such, strain 13 was selected as a robust model for drug combination experiments to evaluate the ability of EIs to restore BDQ susceptibility in *Rv0678* mutants.

Combination assays of BDQ with EPs against the resistant mutant 13 yielded remarkable results. LYO, BER, and PIP emerged as particularly effective, restoring BDQ’s potency to near wild-type levels. These findings are consistent with previous studies demonstrating the inhibitory activity of BER, RES, and PIP against bacterial efflux systems when combined with other antibiotics. For instance, BER targeting NorA in *Staphylococcus aureus*, RES inhibiting AcrAB-TolC in *Escherichia coli*, and PIP modulating MexAB-OprM in *Pseudomonas aeruginosa* (Jin *et al*., 2011; Li & Ge, 2023; Shaheen *et al*., 2019). The MIC of BDQ dropped from 1.563 µM to 0.098 µM in the presence of BER and PIP, representing a 16-fold improvement. LYO increased the potency of BDQ by 16-fold, while RES and LYO-3 had a slight improvement of BDQ MIC by 8-fold and 4-fold, respectively. These effects suggest that efflux is a major contributor to the resistance phenotype and that its inhibition can re-sensitize *Mtb* mutants to BDQ.

Synergistic effects were further confirmed by FICIs. For the BDQ+BER combination, the FICI was 0.19, and BDQ+RES treatment had a FICI of 0.37. An additive effect was noted for the co-treatment of BDQ+PIP, with a FICI of 0.557. Since the individual MICs of LYO and LYO-3 were greater than the highest concentration tested, their FICI values could not be calculated. Nonetheless, the pronounced reduction of BDQ’s MIC observed when in combination with the EIs strongly suggests a synergistic interaction. It is also worth noting that relying solely on FICI values may not fully capture the complexity of synergistic interactions, especially in the context of natural product combinations, which may act through multiple, overlapping mechanisms (Fatsis-Kavalopoulos *et al*., 2024). These findings are consistent with prior studies, which have shown that efflux inhibitors can overcome resistance in other pathogens. The ability of natural products to function as adjunctive agents provides a promising avenue, especially in settings where new drugs are scarce and/or costly (Ramalingam *et al*., 2024).

Beyond simply inhibiting bacterial growth, it was crucial to determine whether drug combinations exhibited bactericidal activity against *Mtb*. The bactericidal potential of the BDQ+LYO combination was assessed using an MBC assay. At specific concentrations such as 0.196 µM BDQ combined with 0.97 µM LYO, the pair achieved a ≥4-log₁₀ reduction in CFU, meeting the standard threshold for bactericidal effect as described by Santos *et al*., 2020. At lower concentrations, the reduction in CFU was less significant and was classified as bacteriostatic. These results indicate that efflux inhibition by LYO not only increases *Mtb’s* susceptibility to BDQ but also enhances its bactericidal activity. This has important therapeutic implications, as bactericidal activity is a key factor in treatment success, especially in preventing disease relapse and limiting transmission potential (Mann *et al*., 2024).

Importantly, LYO and LYO-3 have not, to the best of our knowledge, been investigated as putative efflux inhibitors against *Mtb.* Therefore, their inclusion in this study adds novelty and opens new possibilities for future research in drug development.

## Conclusion

This study suggests that natural product compounds can reverse BDQ resistance by inhibiting efflux activity, thereby restoring the drug’s efficacy. The findings also indicate that BDQ resistance in *Mtb* can arise spontaneously and rapidly under selective drug pressure. Consistent with previous reports, this study confirms that mutations in the *Rv0678* gene appear to represent a central mechanism underlying this resistance, likely through the deregulation and overexpression of efflux pumps.

## Study Limitations

This study provides phenotypic evidence of novel EIs that modulate efflux activity, but several limitations must be acknowledged. Functional efflux assays, transcriptional analysis of MmpS5-MmpL5, genetic manipulation of efflux pumps, and gene expression assays were not performed, leaving the proposed mechanism inferred rather than directly validated. Validation in MDR/XDR *Mtb* strains, medicinal chemistry optimization of EIs, and pharmacokinetic and toxicity profiling were also not undertaken. Importantly, all of these limitations stemmed from inadequate funding and technical constraints rather than a lack of scientific rationale. Future studies should prioritize mechanistic validation, compound optimization, and *in vivo* efficacy to strengthen clinical relevance.

## Acknowledgements

We acknowledge the Science for Africa Foundation for funding this study through the Grand Challenges Africa Program Round 10-2^nd^ Drug Discovery Award (GCA/DD2/Round10/022/002). We greatly appreciate the support we received through a Scientific Discovery Fellowship at Janssen Pharmaceuticals N.V., in partnership with the Johnson & Johnson Foundation, through which this collaboration was initially established. We are further grateful to the Holistic Drug Discovery and Development (H3D) Centre Biology Laboratory at University of Cape Town for their invaluable support in establishing and optimizing the in vitro assays essential to this study.

## Author contributions

Robi Chacha (1,2): Conceptualization, methodology design, data curation, and manuscript drafting. Maes Valerie (3): Formal analysis, creation and validation of BDQ resistant strains, and critical review of the manuscript. Mathew P. Ngugi (2): Investigation, experimental work, and contribution to data interpretation. Edwin K. Murungi (4): Supervision, project administration, guidance on biological assays, and funding acquisition. Dirk A. Lamprecht (5): Technical support for assay development, and manuscript editing. Elizabeth M. Kigondu (6): Supervision, funding acquisition, overall project oversight, and final approval of the manuscript. All authors contributed to the discussion of results, reviewed the manuscript, and approved the final version for submission.

